# A multi-locus association analysis method integrating phenotype and expression data reveals multiple novel associations to flowering time variation in wild-collected *Arabidopsis thaliana*

**DOI:** 10.1101/195446

**Authors:** Yanjun Zan, Örjan Carlborg

## Abstract

When a species adapts to a new habitat, selection for the fitness traits often result in a confounding between genome-wide genotype and adaptive alleles. It is a major statistical challenge to detect such adaptive polymorphisms if the confounding is strong, or the effects of the adaptive alleles are weak. Here, we describe a novel approach to dissect polygenic traits in natural populations. First, candidate adaptive loci are identified by screening for loci that are directly associated to the trait or control the expression of genes known to affect it. Then, the multi-locus genetic architecture is inferred using a backward elimination association analysis across all the candidate loci using an adaptive false-discovery rate based threshold. Effects of population stratification are controlled by corrections for population structure in the pre-screening step and by simultaneously testing all candidate loci in the multi-locus model. We illustrate the method by exploring the polygenic basis of an important adaptive trait, flowering time in *Arabidopsis thaliana*, using public data from the 1,001 genomes project. Our method revealed associations between 33 (29) loci and flowering time at 10 (16)°C in this collection of natural accessions, where standard genome wide association analysis methods detected 5 (3) loci. The 33 (29) loci explained approximately 55 (48)% of the total phenotypic variance of the respective traits. Our work illustrates how the genetic basis of highly polygenic adaptive traits in natural populations can be explored in much greater detail by using new multi-locus mapping approaches taking advantage of prior biological information as well as genome and transcriptome data.

## Introduction

The genome wide association study (GWAS) is a powerful approach to dissect the genetic basis of phenotypic variation in populations, but its application in natural populations is statistically challenging. For example, when populations develop local adaptations to their environment, this often results in a confounding between adaptive alleles and the genome-wide genotype of the local population. This confounding is particularly challenging when adaptations are developed for highly polygenic traits where the selection responses are due to changes in the allele frequencies of standing variants across many loci (Burke, Liti, & Long, 2014; Orozco-terWengel et al., 2012; Teotónio, Chelo, Bradić, Rose, & Long, 2009). The power to detect loci making small contributions to the trait variation is generally low in GWAS due to corrections for multiple testing. The additional burden of corrections also for population structure decreases power further. By making combined use of powerful model populations, and new analytical approaches accounting for the polygenic nature of complex adaptive traits, it is possible to dissect polygenic genetic architecture with greater sensitivity in order to facilitate deeper insights to the genetic basis of adaptive traits (Sheng, Pettersson, Honaker, Siegel, & Carlborg, 2015; Zan et al., 2017). These potent statistical methods have so far not been adapted to analyses of natural populations, but could be useful also for identifying the genetic mechanisms that govern adaptation for polygenic traits in nature.

*Arabidopsis thaliana* is a widely used model species in plant biology. It has colonised a wide range of ecological habitats around the world and the available genotype, expression and phenotype data on many of these ecotypes provides unique opportunities to study the genetic basis of adaptation. Flowering time is one of the most studied adaptive traits in this species as it has an important role in ecological adaptation and also a potential impact on agronomic production in related species. Studies in the laboratory, often on the reference accession *Col-0*, have identified many genes that could influence flowering time and in this way biological pathways controlling, for example, photoperiod (El-Assal, Alonso-Blanco, Peeters, Raz, & Koornneef, 2001; Filiault et al., 2008), vernalization (Li et al., 2014; Shindo et al., 2005) and plant hormone signalling (Sharma et al., 2016) have been identified. Studies of flowering time variation in nature includes both QTL mapping in crosses between divergent accessions and genome wide association (GWA) studies in collections of wild accessions (Alonso-Blanco et al., 2016; Atwell et al., 2010; Brachi et al., 2010; Salomé et al., 2011). The molecular studies suggest that hundreds of genes could potentially influence flowering time whereas the studies of natural populations have only been able to associate genetic polymorphisms in a handful of them with the large world wide flowering time variation. Hence, even though flowering time is potentially highly polygenetic also in nature, the burden of multiple-testing and population-stratification correction in GWAS analyses of natural associations is too high to identify loci that are either rare, have smaller effects or are confounded with population structure even in collections of >1,000 inbred accessions (Alonso-Blanco et al., 2016).

Here, we describe an analytical approach to dissect the genetic basis of polygenic adaptive traits in natural populations. It is used to dissect flowering time variation in a publicly available dataset on >1,000 wild-collected *A. thaliana* accessions from the 1,001-genomes project (Alonso-Blanco et al., 2016). By utilizing prior information on known flowering time genes and flowering time associated loci in *A. thaliana* and integrating it in our analysis of genotype, expression and trait variation data from the 1,001-genomes project, we could reveal a highly polygenic basis of the worldwide variation in flowering time.

## Results

We explore the polygenic basis of flowering-time in the latest release of the 1,001 genomes *A. thaliana* dataset. First loci with direct associations to flowering time, or loci that regulate the expression of known flowering time genes, were identified. Then a final set of loci with independent associations to flowering time was obtained using a multi-locus association analysis approach that was originally developed to dissect polygenic traits in experimental populations (Brandt, Ahsan, Honaker, & Siegel, 2017; Lillie et al., 2017; Sheng et al., 2015; Zan et al., 2017).

### A genome wide association analysis to detect new loci associated with flowering time

A publicly available dataset with genotypes and phenotypes measured on a collection of 1,004 natural *Arabidopsis thaliana* accessions was reanalysed. Two flowering time phenotypes measured under greenhouse conditions, flowering time at 10°C (FT10) and 16°C (FT16), were used in a previous genome wide association study by (Alonso-Blanco et al., 2016). Here, we reanalyse these phenotypes and describe the results for FT10 in more detail. A more schematic summary of the results is provided for FT16.

Five loci were associated with FT10 in (Alonso-Blanco et al., 2016) using a mixed model analysis accounting for population structure (Figure 1A). We here explore the genetic architecture of FT10 beyond those loci. First, the five earlier reported loci were included as co-factors in genetic model. Then, the genome was scanned iteratively to identify additional loci directly associated with FT10. In this scan, we eased the burden of multiple-testing correction by only correcting for tests performed on markers in LD < 0.95, resulting in a selection threshold for terminating the forward selection scan of -log_10_p =7.33. Two additional loci were associated with FT10 at this significance level after correction for population structure via a mixed model analysis (Figure 1B). One of these associations (peak on chromosome 4, 17,282,047bp) overlapped with the confidence interval of an earlier reported flowering time QTL in *A. thaliana* recombination inbred lines ((Salomé et al., 2011); Table 1). In addition to these loci, a third locus (peak on chromosome 4, 10,999,188bp; Figure 1C) had a p-value near the selection threshold (-log_10_ p = 7.0). As it overlapped with the confidence interval of an earlier reported flowering time QTL ((Salomé et al., 2011); Table 1) and only 3.8 kb upstream of the known flowering time gene *TSF* (Twin sister of FT, AT4G20370 (Ando et al., 2013)), it was retained for the multi-locus analysis.

**Figure 1.**
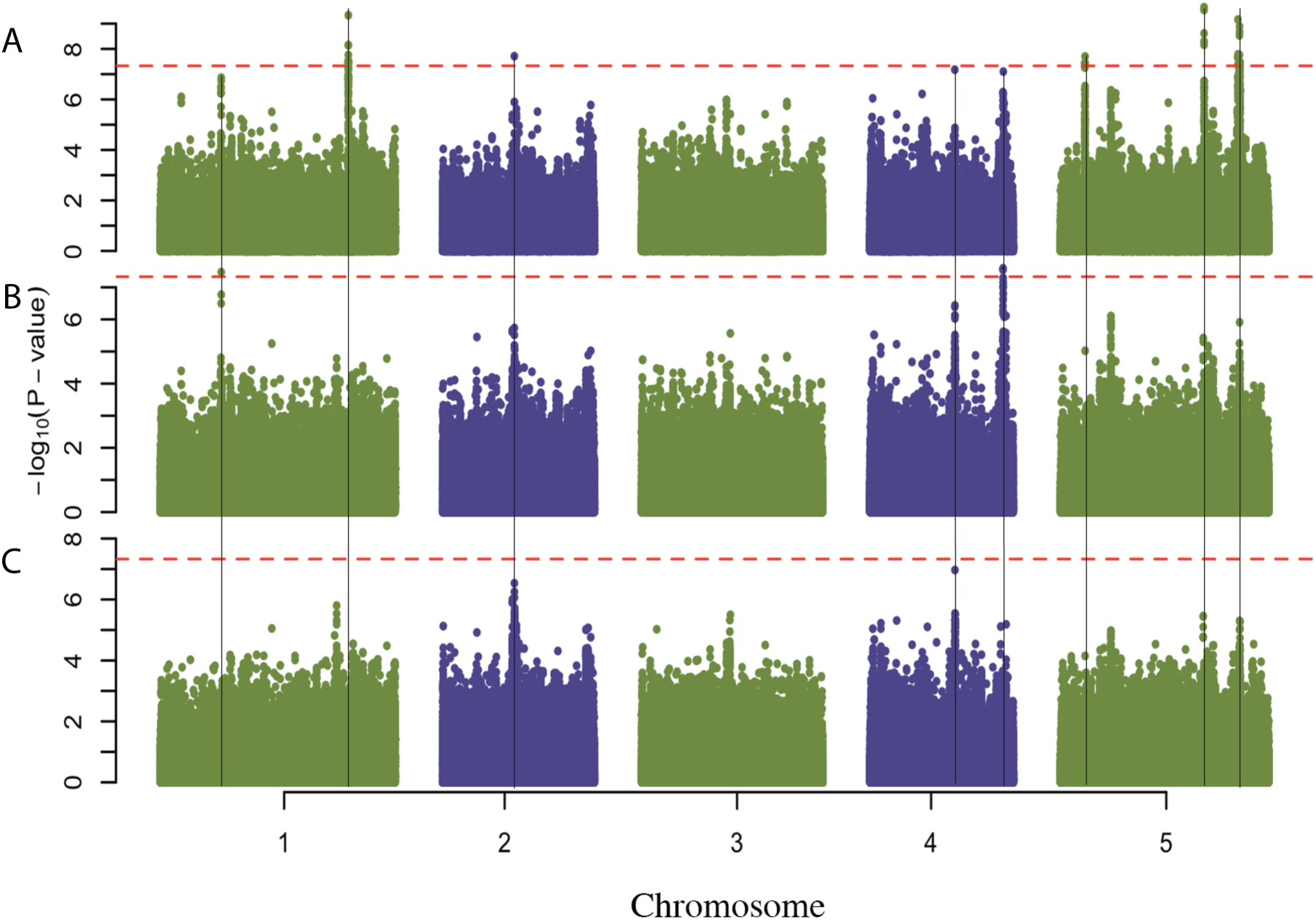
*Intermediary Manhattan plots from the forward selection genome scans to detect loci associated with flowering time at 10°C (FT10) in the 1001-genomes* A. thaliana *collection* (Alonso-Blanco et al., 2016). ***(A)*** *GWA results for FT10 without genetic covariates in the model replicates the 5 loci in* (Alonso-Blanco et al., 2016). ***(B)*** *GWA results for FT10 with the 5 significant loci in (A) as covariates identifies the two additional FT10 loci passing the selection threshold in the forward selection GWA scan.* ***(C)*** *Final GWA results for FT10 where all 7 loci from (A) and (B) passing the forward selection termination threshold were included as covariates. The 8*^*th*^ *selected locus was near the termination threshold (-log*_*10*_*p = 7.0 vs 7.3 for the selection threshold indicated as a dashed red horizontal line). The loci selected for the multi-locus analysis are highlighted with vertical lines.*

**Table 1.** *Eight loci selected for the multi-locus analysis based on direct associations to flowering time at 10°C.*

| Chr | Pos<sup>1</sup> | Late flowering allele<sup>2</sup> | Freq<sup>3</sup> | a<sup>4</sup> | p-value<sup>5</sup> | Var<sup>6</sup> |
| --- | --- | --- | --- | --- | --- | --- |
| 1 | 7,885,760 | T | 0.25 | 2.2±0.4 | 1.8×10<sup>-6</sup> | 5.5% |
| 1 | 24,339,560 | G | 0.56 | 2.5±0.4 | 2.2×10<sup>-9</sup> | 8.0% |
| 2 | 9,312,986 | A | 0.09 | 2.9±0.6 | 7.1×10<sup>-7</sup> | 4.2% |
| 4 | 10,999,188<sup>#</sup> | G | 0.87 | 4.5±0.7 | 6.0×10<sup>-9</sup> | 2.5% |
| 4 | 17,282,047<sup>#</sup> | T | 0.05 | 5.1±0.9 | 1.0×10<sup>-7</sup> | 1.4% |
| 5 | 3,178,338 | A | 0.65 | 2.8±0.5 | 7.2×10<sup>-8</sup> | 6.3% |
| 5 | 18,590,327 | A | 0.53 | 2.7±0.5 | 8.0×10<sup>-8</sup> | 11.4% |
| 5 | 22,979,827 | T | 0.04 | 5.6±0.9 | 1.8×10<sup>-9</sup> | 7.4% |

<sup>1</sup>Location of peak SNP in bp; <sup>2</sup>SNP allele associated with delayed flowering at 10°C; <sup>3</sup>Frequency of SNP allele delaying flowering at 10°C; <sup>4</sup>Estimated additive effect of the locus in days ± standard error; <sup>5</sup>p-value for association and <sup>6</sup>additive genetic variance explained by the locus, all obtained from a mixed model including all eight selected loci and the kinship matrix to account for population structure; <sup>#</sup>SNP location overlaps with 95% QTL confidence interval of (Salomé et al., 2011).

### Loci associated with expression variation in earlier reported flowering time genes

Molecular studies in the reference *Arabidopsis thaliana* accession *Col-0* have revealed many genes that are directly or indirectly involved in the regulation of flowering time (Brachi et al., 2010). The 1,001 genomes data was screened for polymorphisms associated with expression variation in 282 known flowering time genes (Brachi et al., 2010) using a mixed model based expression QTL (eQTL) mapping analysis accounting for population structure. In total, 150 eQTL regulated the expression of 123 flowering time genes at genome wide significance (Table S1). One locus on chromosome 5 (18,590,327bp) regulated expression of the well-studied flowering time gene *DOG1* (*AT5G45830*). Another locus on chromosome 5 (22,603,100bp) acted as a trans eQTL hotspot regulating the expression of 33 flowering time genes (Figure 2; Table S2). Among the 33 genes regulated by this trans eQTL, 19 have a TAIR10 annotation/GO term related to flowering time via light mediated developmental, or photoperiod control, pathways (Table S3). The LD block around the top associated SNP (r^2^ > 0.5) spans 12kb on chromosome 5 including 5 genes (*AT5G55830*, *AT5G55835*, *AT5G55840*, *AT5G55850* and *AT5G55855*). An obvious candidate among those is *AT5G55835* as it is known to be expressed as a microRNA regulating many developmental processes, including changes in the vegetative phase, and coping with abiotic stress (Lei, Lin, & An, 2016; Wang, Czech, & Weigel, 2009; Yang, Conway, & Poethig, 2011; Yu et al., 2015).

**Figure 2.**
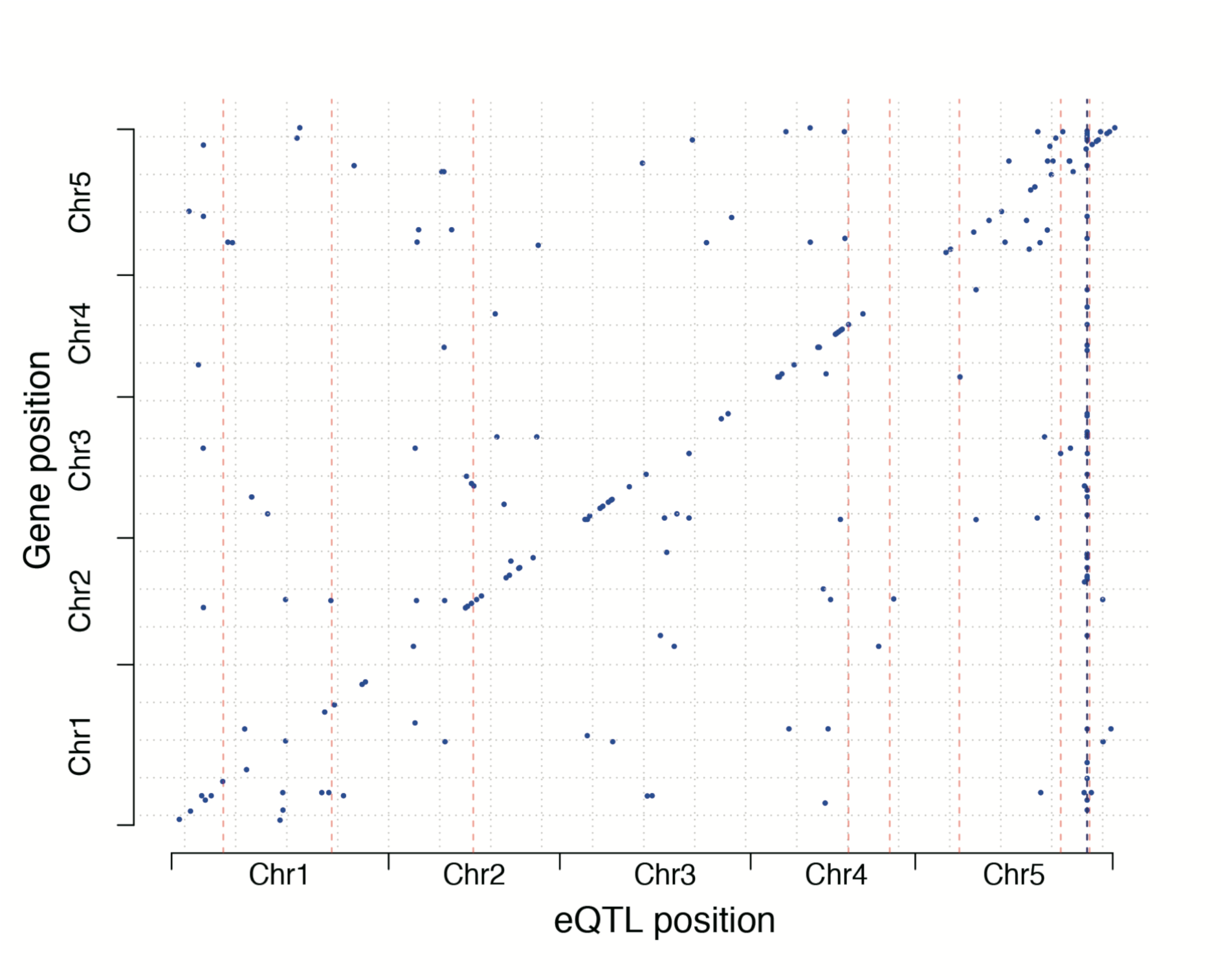
*A trans-eQTL hotspot on chromosome 5 regulates the expression of 33 flowering time genes in the 1,001 genomes collection. The genomic locations of 123 known flowering time genes* (Brachi et al., 2010)*, whose expression are regulated by eQTL segregating in the 1,001 genomes collection, are given on the y-axis. The locations of the eQTL regulating the expression of the flowering time genes are on the x-axis. The eight loci with direct associations to FT10 (Table 1) are highlighted as dashed pink lines and the trans-eQTL hotspot as a dashed black line.*

### Multi-locus analysis of loci Multi-locus analysis associated with FT10 or the expression of flowering time genes

A backward-elimination, model-selection analysis with an adaptive False Discovery Rate (FDR) criterion (Brandt et al., 2017; Lillie et al., 2017; Sheng et al., 2015; Zan et al., 2017) was used to identify a polygenic model of loci associated with FT10 in the 1,001 genomes collection. The input to the analysis were the loci with direct associations with FT10 in our forward selection genome-wide association analysis (Table 1) and the eQTL regulating the expression of known flowering time genes (Table S1). At 15% FDR, 25 of the eQTL and all 8 FT10 associated loci were retained in the final model (Figure 3; Table S3). At a 15% FDR, we expect 28 of the 33 loci to be true associations, illustrating the ability of the method to reveal the highly polygenic architecture of FT10 in this population.

**Figure 3.**
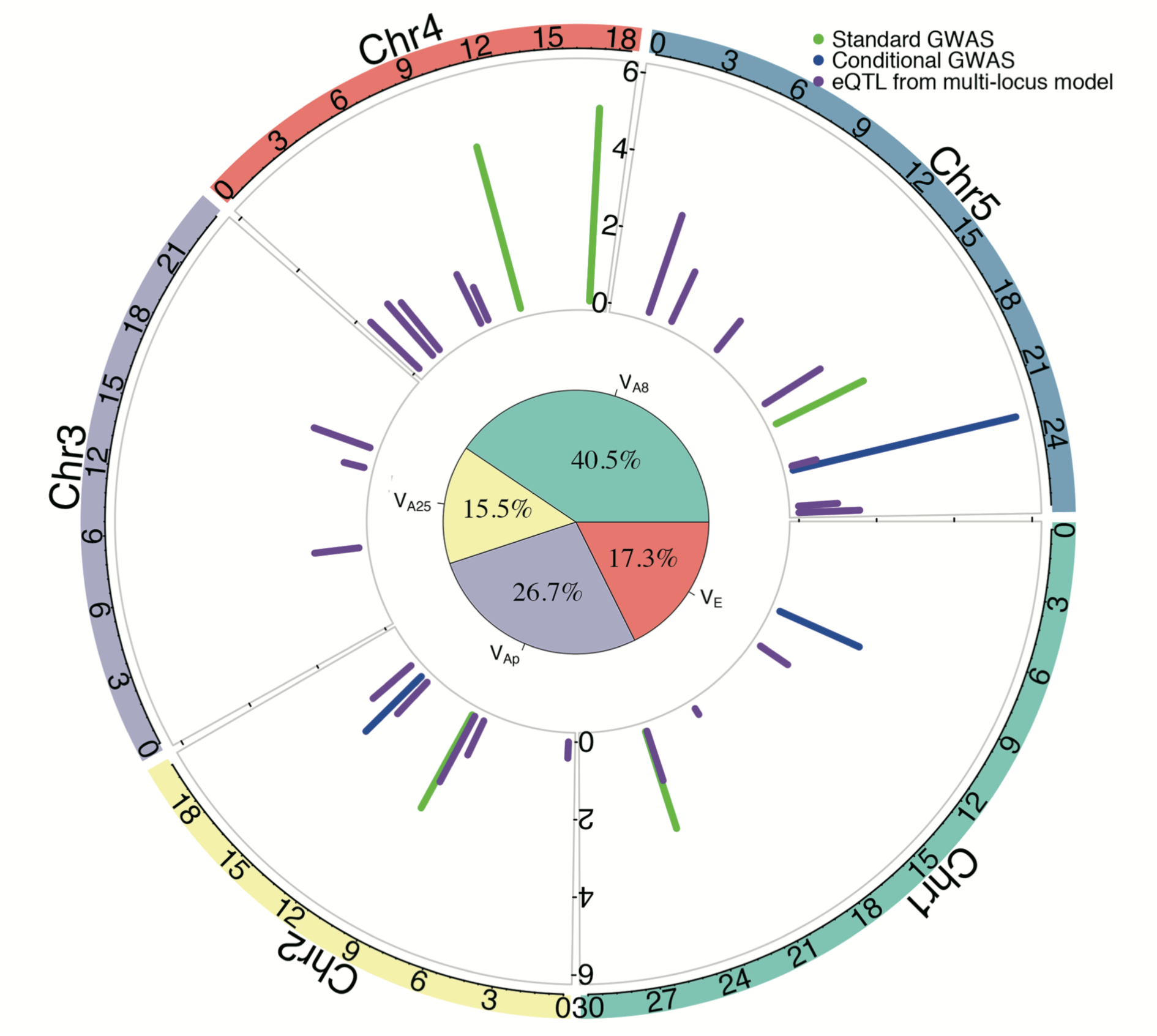
*An illustration of the polygenic architecture of flowering time at 10°C in the green house of the 1,001 genomes Arabidopsis thaliana collection. The five chromosomes are illustrated as coloured bars around the circle perimeter. The green lines illustrate the additive effects of the 5 QTL associated with FT10 in* (Alonso-Blanco et al., 2016) *, the blue lines the additive effects of the 3 additional QTL selected in our forward-selection GWA and the purple lines the additive effects of the 25 eQTL associated with FT10 in the multi-locus analysis. The pie chart in the middle shows the proportion of phenotypic variance explained by the 8 loci with direct associations to FT10 (V*_*A8*_*), the 25 eQTL (V*_*A25*_*), the additional polygenetic effect explained by the kinship matrix (V*_*Ap*_*) and the residual environmental effects (V*_*E*_).

The additive genetic effects of the associated loci range from 1.4 to 7.6 days with median of 2.5 (Figure 4A). The proportion of the total genetic variance for FT10 that could be explained by fitting the Identity by State genome wide kinship matrix is 82%. The 33 FT10 associated loci together explain 67% of the genetic variance captured by the kinship matrix, with 3/4 of this contributed by the 13 loci with the largest effects (Figure 4B). The additive variance contributed by the individual locus with the largest effect (chromosome 5 at 18,590,327bp) is 11% of the total genetic variance in flowering time in this population.

**Figure 4.**
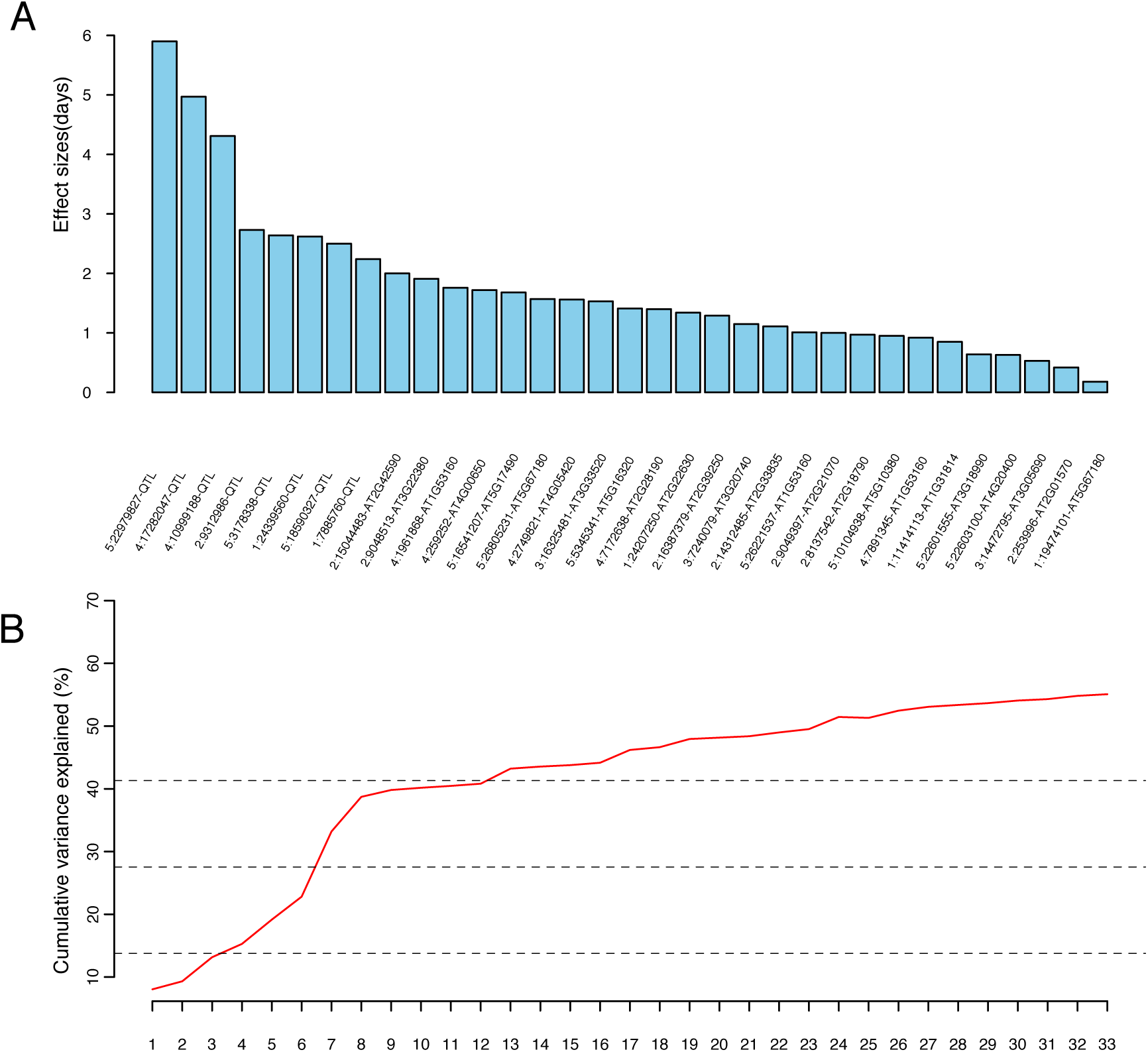
***(A)*** *Histogram illustrating the additive effects for the 33 loci associated with FT10 in our multi locus analysis at a 15% FDR.* ***(B)*** *Cumulative variance explained by the 33 loci (read line). These estimates were calculated by adding one locus at a time in the model in the order determined by the effect sizes in (A). The 25%, 50% and 75% quartiles of the total variance explained by the loci are highlighted using dashed black lines.*

### Large overlap between loci associated with flowering times at 10°C and 16°C

To explore the overlap between the polygenic architectures of FT10 and FT16, we performed an identical three-step analysis also for FT16. The two phenotypes have a high phenotypic correlation (Pearson correlation=0.88, p-value < 2.2×10^-16^), but the standard GWAS analysis only detects 3 loci for FT16 at a genome wide significance level. Our conditional forward selection GWA analysis for FT16 failed to reveal any additional loci directly associated with this trait (Figure S1, Table S1). All the three loci associated with FT16 map within 20kb of the peak SNPs associated with FT10 (Table S4). The multi-locus analysis across the 150 eQTL regulating expression of flowering time genes and the three FT16 associated loci identified 29 loci associated with FT16 at 15% FDR. In total, 15 of the 26 eQTL overlapped with the 25 eQTL associated with FT10. (Table S5). The three loci with direct associations to FT16 together explain 19% of the genetic variance captured by the genomic kinship matrix, and the 26 eQTL detected in the multi-locus analysis explain an additional 29%.

## Discussion

Here, we describe a multi-locus method to dissect polygenic genetic architectures of adaptive traits in populations of wild-collected *A. thaliana* accessions. It integrates prior knowledge about genes affecting the adaptive trait from earlier functional studies with experimental genome and transcriptome data on a large collection of natural accessions from the 1,001 genomes project (Alonso-Blanco et al., 2016; Kawakatsu et al., 2016) to facilitate mapping of alleles with small effects on the trait. In our study, we test the method on a well studied adaptive trait – flowering time – where among 282 known flowering time genes (Brachi et al., 2010), the expression of 123 are controlled by polymorphisms segregating in the 1,001 genomes collection. In a multi-locus analysis (Abramovich, Benjamini, Donoho, & Johnstone, 2006; Brandt et al., 2017; Lillie et al., 2017; Sheng et al., 2015) where we simultaneously test the effect on flowering time of loci with GWA associations to flowering time and eQTL, 33 loci were shown to be associated with variance in flowering time at 10°C (FT10). For flowering time at 16°C (FT16), 29 loci were detected using the same approach. Of these, 8 (3) were detected in the GWA analysis and 25 (26) in the eQTL analysis for FT10 (FT16), respectively, at a 15% FDR threshold. Together these results illustrate the value of using a more sensitive analysis approach to infer the polygenic architecture of complex adaptive traits in natural populations. The results illustrate that many loci contribute small effects to the phenotypic variation in the population that would otherwise remain undiscovered. Overall, the genetic architecture thus contains a relatively small number of loci with large effects, or loci with intermediate effects but high MAF, that can be detected in a standard genome-wide association analysis. In addition to this, our analysis reveals a larger number of loci that together explain a relative large amount of the total genetic variation. The multi-locus analysis thus provides a deeper insight to this polygenic adaptive trait to facilitate further in-depth studies about how the loci together contribute to local and global adaptation in the world wide population.

In total, eight (three) large effect loci were detected in the GWA analyses for FT10 (FT16). Together they contributed 40 (19)% of the genetic variance for FT10 (FT16) in this population. This finding is consistent with earlier reports in *Arabidopsis thaliana* that major loci often make important contributions to adaptive traits (Brachi et al., 2015; Forsberg et al., 2015; Rus et al., 2006; Shen et al., 2014; Shindo et al., 2005). Here, we can also show that in addition to the major loci, many others also contribute to the variation in this adaptive phenotype. These are difficult do detect in the GWA, either due to them contributing with individually small effects, or by them having a low minor allele frequency. Together, however, they contribute a considerable fraction of the flowering time variation in the 1,001 genomes collection and especially so for FT16. This finding suggests that polygenetic adaptation, in addition to major alleles, is likely to make an important contribution to adaptation in *A. thaliana*. This is consistent with earlier experimental findings on studies of artificial selection responses, where it has been shown that selection on many loci can lead to extreme adaptations in, for example, seed protein and kernel oil in maize (Laurie et al., 2004; Lucas, Zhao, Schneerman, & Moose, 2013), and body weight in mice (Allan, Eisen, & Pomp, 2005) and chicken (Besnier et al., 2011; Johansson, Pettersson, Siegel, & Carlborg, 2010; Sheng et al., 2015; Wahlberg et al., 2009).

In conclusion, our study demonstrates the value of integrating prior information on the genetic basis of a trait with multiple types of experimental data to dissect polygenic adaptive traits. Using information on known flowering time genes, and experimental data on gene expression, flowering time variation and genome wide SNP variation, a highly polygenic basis was revealed for this trait in the 1,001 genomes collection. The results suggest that a combination of major effect alleles and polygenic adaptation has been important for flowering time adaptation in *A. thaliana*.

## Materials and methods

### Data

All genotype, phenotype and transcriptome data are publicly available within the *Arabidopsis thaliana* 1001 genomes project (Alonso-Blanco et al., 2016; Kawakatsu et al., 2016). The imputed whole genome SNP data matrix was downloaded from http://1001genomes.org/data/GMIMPI/releases/v3.1/SNP_matrix_imputed_hdf5/1001_SNP_MATRIX.tar.gz. We filtered for minor allele frequency and only retained loci with MAF>0.03. SNP markers were pruned to remove loci in pairwise LD of r^2^ > 0.99. In total, 1,396,438 SNPs on 1,004 individuals remained. Flowering times measured at 10°C and 16°C in the green house were downloaded from (Consortium, 2016a, 2016b). In total 728 transcriptomes were downloaded from NCBI (GSE80744), and 648 of these were used in the expression QTL mapping analysis as the remaining ones had no matching genotypes in the

### Genome wide association analysis for flowering time

Previous genome wide association (GWA) analysis in this dataset reported 5 genome-wide significant associations for flowering time at 10°C (16°C) using a mixed model analysis to account for population structure (Alonso-Blanco et al., 2016). We here extended these analyses for these traits by first including the already detected loci as cofactors in the analysis model and then performing several iterative GWA analyses where new loci were added to the model if they passed the multiple-testing corrected genome-wide significance threshold. The termination threshold for the forward selection scans was calculated using a Bonferroni correction for the number of SNPs with r^2^ < 0.95, resulting in a threshold of –log_10_p = 7.33. The model used in the GWA analyses was:

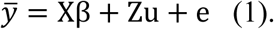

Where 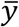 is the mean flowering time at 10°C (16°C), *X* the design matrix with the number of columns equal to the number of already selected SNPs plus one for the SNP tested for inclusion (coded as 0, 2 for minor-allele homozygous, major-allele homozygous genotypes, respectively), *β* is a vector of the effects of substituting two alleles at the corresponding SNPs in *X*, *Z* is the design matrix obtained from a Cholesky decomposition of the *G* (identity by state – IBS) kinship matrix estimated from the whole-genome SNP data with the *ibs* function in the GenABEL package (Aulchenko, Ripke et al. 2007). The *Z* matrix satisfies *ZZ*′ = *G*, thus, the random effect vector *u* will be normally distributed, 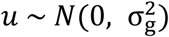, *e* is the normally distributed residual variance with 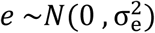, The analyses used the *polygenic* and *mmscore* functions in GenABEL package (Aulchenko, Ripke et al. 2007).

### Expression QTL mapping targeting genes in flowering time related pathways

A list of 282 flowering time genes was obtained from (Brachi et al., 2010). From the available transcriptome sequence data in the 1,001 genomes project, the expression levels were extracted for those genes as described in (Zan, Shen, Forsberg, & Carlborg, 2016). Whole-genome eQTL mapping was then performed for these genes across the pruned SNP set (r^2^ > 0.99) described above. In the analysis, model 1 was used but with a slightly altered design matrix ***X***. Here, ***X*** was only a column vector containing the genotype of the tested SNP, coded as 0, 2 for minor-allele homozygous, major-allele homozygous genotypes. As in the GWA, a Bonferroni corrected significance threshold was used and calculated based on the number of SNPs in the dataset with r^2^ < 0.95, resulting in a multiple-testing corrected threshold of –log_10_p=7.33.

### Multi-locus backward elimination association analysis

We have earlier developed a multi locus approach to explore the genetic architecture of highly polygenic traits and a detailed description of the approach is available in (Sheng *et al.* 2015; Brandt *et al.* 2017; Lillie *et al.* 2017). In short, the method was designed to study traits where earlier data suggest them to be polygenic in the analysed population. In such cases, standard association analysis approaches to detect individual loci using stringent multiple testing corrected significance thresholds will have low power to infer the true genetic architecture of the traits. This as many of loci contributing to a polygenic trait will make small individual contributions to the genetic variance. By implementing a multi-locus mapping approach in a backward elimination model selection framework, it is possible to simultaneously account for the joint genetic effects of many loci and identify those that together contribute to the trait variance. Not known in advance is how many loci will contribute to the trait and therefore the number of loci in the final model could vary substantially. To account for this our analysis was implemented with an adaptive model selection criterion to control the False Discovery Rate (FDR) in the final model (Abramovich et al., 2006; Gavrilov, Benjamini, & Sarkar, 2009). By using a 15 % FDR as the termination criterion of the analysis, most of the selected loci are expected to be true associations to the trait. This makes it possible to gain overall insights to the polygenic basis of the studied trait from the mapped loci, but as the significance of individual loci is not tested it is not suitable to use the analysis to make conclusions about these.

The multi-locus analysis was here extended to identify a multi-locus model across loci with direct associations to flowering time and loci that might influence flowering time via their transcriptional regulation of known flowering genes. The major focus of the multi locus analysis was here to evaluate the joint contribution of all these loci to flowering time variation. In particular, the analysis would reveal if using information on flowering gene eQTL was important for dissecting the genetic basis of flowering time variation in this dataset.

The first step of the multi-locus analysis was a pre screening, where the 150 significant eQTL markers were divided into 3 sets with approximately 50 markers each. The markers were divided to include markers on the same chromosomes. This step was used since a backward elimination analysis across >100 markers would be instable due to an over parameterized linear model. For each of pre screening sets, a backward-elimination analysis was performed with termination based on the adaptive FDR criterion. Markers reaching the 15% FDR in either of these analyses were kept for joint analyses.

The second step of the analysis was a full analysis, where all markers selected in the pre-screening of the eQTL loci and the loci with direct associations to flowering time were analysed jointly. The end result was a set of loci that together contributes to flowering time variation in this population at a 15% FDR. All analyses were implemented in custom scripts in the statistical software R (R Core Team 2015) and these are available in the R package *BE* on github (https://github.com/yanjunzan/BE). The genetic effects of those 33 loci, modelled as the difference between the two homozygous genotypes, and the corresponding p value of all associated loci were subsequently estimated using a mixed model fitted using the *hglm* function in R package *hglm* (Rönnegård, Shen, & Alam, 2010).

### Estimation of the variance explained by the mapped loci

Every accession in the dataset was grown in replicates and the average flowering time provided in (Alonso-Blanco et al., 2016; Kawakatsu et al., 2016). To estimate proportion of the total variance in mean flowering times explained by the kinship between the accessions, we fitted a mixed model 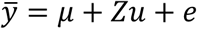 to the data. Here, 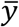 is the mean flowering time per accession and *ZZ*^T^ = *G*, where *G* is the genomic kinship matrix defined above. The intra-class correlation 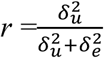 is here the amount of variance in 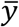 explained by the kinship between the accessions. Under the assumption that the replicated measurements of flowering time within each accession have removed all environmental variance, the total variance represents the genetic variance for this phenotype. The amount of variance explained by the kinship would then equal the proportion of genetic variance that is additive genetic variance. In reality, the number of replicates is likely too low to remove all environmental noise, making the total variance an overestimate of the total genetic variance, and the variance explained by kinship a lower bound of the true proportion of additive variance.

The fraction of the genetic variance explained by a particular set of mapped loci (represented by one marker each) was estimated as 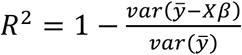 where 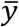 is the mean flowering time per accession and *X* the genotype matrix for the markers in the set fitted as a fixed effect. We here assume that 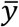 contains no environmental variance, but as it is likely to the obtained estimate will instead be the lower bound of the proportion of the genetic variance explained by the additive effects of the set of markers, in the same way as for the estimate of the variance by the kinship described above. When comparing the variances explained by different sets of loci, we first estimated the variance explained by all loci (V_full_), then the variance of the reduced set (V_red_) where the markers whose contribution was to be estimated are excluded, to then estimate the variance they contributed as Vf_ull_ - V_red_.

### Visualization of the results

Figure 3 was made using the R package *Circlize* (Gu, Gu, Eils, Schlesner, & Brors, 2014).

## Acknowledgements

This work was supported by the Swedish Research Council for Environment, Agricultural Sciences and Spatial Planning (Formas grant ID 2013-450 to ÖC).

## Author contributions

ÖC and YZ initiated the study, designed the project and the statistical analyses; YZ wrote the analysis scripts and performed the data analyses. ÖC and YZ summarized the results and wrote the manuscript.

## Disclosure declaration

The authors declare no competing interest.

## Supplementary material

Supplementary Tables S1-S5 and Supplementary Figure S1 are provided in the file Supplementary_data.pdf

